# ApoFLARE: a luminescent reporter for direct quantification of APOBEC3A editing activity

**DOI:** 10.64898/2026.03.13.710312

**Authors:** Maria Vittoria Di Marco, Braeden L. Butler, Christopher T. Eggers, Aaron N. Hata

## Abstract

APOBEC-mediated cytidine deamination is a major endogenous source of mutagenesis in human cancers and has been linked to tumor evolution, clonal diversification and therapeutic resistance. Among the APOBEC family, APOBEC3A (A3A) is a potent and inducible cytidine deaminase, with dynamic and context-dependent activation. Most approaches for studying the role of A3A in cancer infer A3A activity indirectly via its expression level or retrospective mutational signatures, or through molecular assays that are limited to endpoint measurements and do not readily allow longitudinal interrogation of A3A editing dynamics. Therefore, quantifying the timing, persistence, and cellular heterogeneity of A3A activity remains challenging. Here, we describe ApoFLARE, a genetically encoded reporter that converts A3A-mediated cytidine deamination into a quantitative luminescent signal in living cells. ApoFLARE allows for scalable, ratiometric measurement of editing activity and enables time-resolved analysis of editing kinetics. Reporter activation is selectively dependent on A3A catalytic function and was absent in A3A-deficient, but not A3B-deficient cells. Under stress and targeted therapy conditions, reporter activity correlated with endogenous RNA editing measured by digital droplet PCR, including contexts in which catalytic activity persisted beyond transient A3A transcript induction. Thus, ApoFLARE offers a scalable platform to investigate the regulation, kinetics, and heterogeneity of A3A editing.

**GRAPHICAL ABSTRACT:** 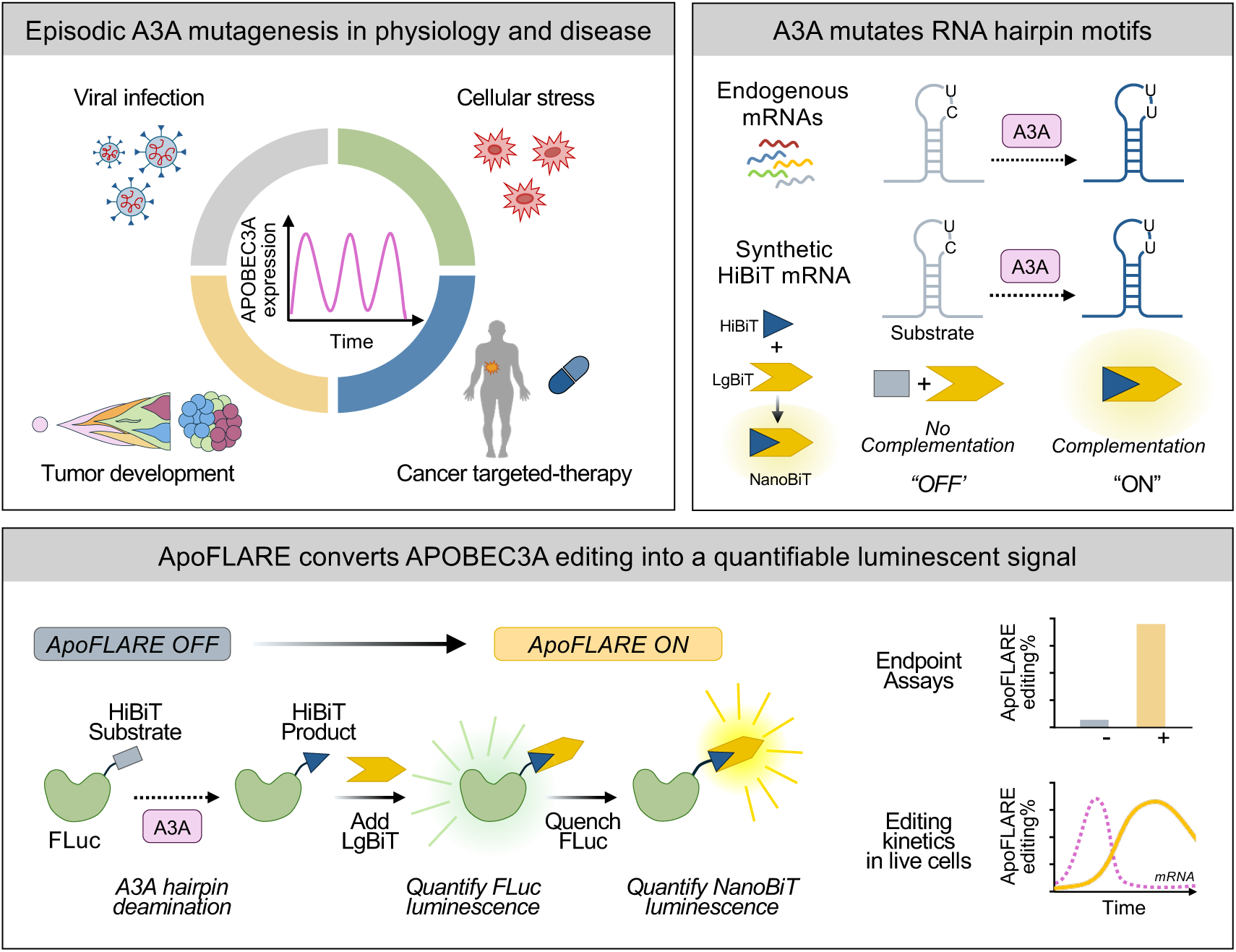

## INTRODUCTION

Cytidine deamination is a fundamental nucleic acid-modifying reaction that contributes to sequence diversification across biological systems. In mammalian cells, members of the APOBEC cytidine deaminase family catalyze the conversion of cytidine to uridine in single-stranded nucleic acids, with individual enzymes exhibiting distinct substrate preferences, regulatory mechanisms, and biological functions (1, 2). While APOBEC enzymes play essential roles in innate immunity and antiviral defense, their dysregulation has emerged as a major endogenous source of genomic instability in cancer, shaping mutational landscapes and influencing tumor evolution and drug resistance (3–9).

Among the APOBEC family, APOBEC3A (A3A) and APOBEC3B (A3B) are thought to be the major contributors to C>G and C>T substitutions at TpC motifs that comprise APOBEC mutational signatures in cancer (Sanger 2 and 13) (4, 5). Although the catalytic domains of A3A and A3B share significant sequence similarity, the two enzymes differ markedly in regulation and activation dynamics (10). Whereas A3B displays a more constitutive nuclear presence, A3A is tightly regulated and inducible, becoming activated in response to inflammatory cues and targeted therapy (11). Additionally, A3A is distinguished by higher intrinsic deaminase activity compared with A3B (12). These features confer substantial mutagenic potential and position A3A as a powerful, context-dependent driver of cytidine deamination during oncogenic perturbation and therapeutic stress.

Despite increasing recognition of A3A as a driver of mutagenesis, methods to directly measure A3A editing remain limited. Many studies have inferred APOBEC activity from transcript abundance and quantification of APOBEC-associated mutational signatures (13, 14). While highly informative for reconstructing historical mutagenic processes in cancer genomes, mutational signatures are inherently retrospective and do not report ongoing or context-specific enzymatic activity (4, 5, 15). Moreover, due to the episodic nature of A3A mutagenesis, gene expression levels at one point in time may not reflect expression levels at the time of active mutagenesis. Additionally, A3A protein and editing can persist even as transcript levels decline, highlighting a potential lack of correlation between mRNA expression and functional editing output (4, 15).

Because A3A and A3B generate similar mutational signatures yet exhibit distinct regulatory control, experimental tools that selectively report A3A-specific editing activity are essential for defining enzyme-specific contributions to tumor evolution. Recently, several approaches have been reported for selectively quantifying A3A editing that take advantage of subtle differences in the preferences of the two enzymes for specific local sequence features. Specifically, biochemical and cellular studies have demonstrated that A3A efficiently deaminates structured single-stranded substrates that can adopt stem-loop hairpin configurations (15–17). A digital droplet PCR (ddPCR) assay that quantifies A3A editing at a hairpin loop motif in the DDOST gene has become a benchmark for functional quantification of A3A editing (15). Using a similar conceptual approach, we developed a computational approach for quantifying A3A editing across ~2000 hairpin motifs across the transcriptome (8). One limitation of these and other assays, however, is that they are limited to endpoint analyses and cannot be used for longitudinal or live-cell interrogation (18, 19). Consequently, key questions, including the kinetics of A3A editing, subpopulation heterogeneity, and their impact on clonal dynamics in response to oncogenic or therapeutic stress, remain difficult to address using existing approaches.

Here, we describe ApoFLARE (<u>APO</u>BEC3A <u>F</u>unctional <u>L</u>uminescent h<u>A</u>irpin <u>RE</u>porter), a genetically encoded, HiBiT-based, dual-luciferase reporter that converts A3A-mediated cytidine deamination into a quantitative bioluminescent signal. HiBiT is an 11-amino-acid peptide tag that binds with high affinity to LgBiT, the large subunit used in NanoLuc® Binary Technology (NanoBiT®), to form the bright, luminescent NanoBiT® enzyme. ApoFLARE exploits A3A structural substrate preference by embedding an engineered hairpin site within the HiBiT-encoding RNA that disrupts complementation with the LgBiT subunit. Upon A3A-mediated C>U deamination and subsequent codon change, complementation is restored to generate functional NanoBiT® luciferase. Fusing HiBiT to firefly luciferase enables ratiometric endpoint quantification as well as longitudinal live-cell monitoring of editing activity. Thus ApoFLARE achieves selective and quantitative measurement of A3A catalytic activity in living cells, providing a scalable assay for investigating the temporal and contextual regulation of APOBEC3A editing.

## MATERIALS AND METHODS

### Biological resources

All cell lines used in this study are listed in Supplementary Table 1. PC9, H3255, H3122, and 293T cells were obtained from the MGH Center for Molecular Therapeutics. HeLa cells were purchased from ATCC. Patient-derived cell lines were established in our laboratory from core biopsy or pleural effusion samples as previously described (24, 25).

All patients provided informed consent to participate in a Dana-Farber-Harvard Cancer Center Institutional Review Board-approved protocol, authorizing research on their samples. All cell lines were maintained in RPMI medium (Life Technologies) supplemented with 10% fetal bovine serum (FBS, Life Technologies) except for 293T and HeLa cells, which were maintained in DMEM medium (Life Technologies) with 10% FBS, and doxycycline-inducible cells, which were maintained in RPMI medium (Life Technologies) with 10% Tet system-approved FBS (Gibco). All experiments were performed in RPMI with 10% FBS. All cells were routinely tested and verified as being free of mycoplasma contamination.

A3A Reporter constructs were designed based on a CMV-driven expression construct for firefly luciferase (luc2) fused to HiBiT at its C-terminus with an 8-residue Gly/Ser linker. Six pairs of reporter constructs were generated by altering the DNA sequence for HiBiT and/or the linker to represent a putative substrate sequence for A3A and its matching product sequence. The new DNA constructs were generated by GenScript by replacing the linker/HiBiT sequence in the original expression construct with the new reporter sequences. pInducer20_A3A^WT^, pInducer20_A3A^E72A^ and pInducer20_A3B^WT^ plasmids were a gift from Remi Buisson (16, 26). pLV-EF1a-IRES-Puro and pLV-EF1a-IRES-Blast were obtained from Addgene (Addgene plasmid # 85132; http://n2t.net/addgene:85132; RRID:Addgene_85132 and Addgene plasmid # 85133; http://n2t.net/addgene:85133; RRID:Addgene_85133) (27).

### Reagents

For immunoblotting, the antibodies used in this study are listed in Supplementary Table 2. For cell culture studies, osimertinib (Osi, third-generation EGFR inhibitor) and lorlatinib (Lor, third-generation ALK inhibitor) were dissolved in DMSO to a final concentration of 10 mmol l^−1^ and stored at −20 °C. Doxycycline (Dox) was dissolved in DMSO to a final concentration of 1 mg ml^−1^ and stored at −20 °C. Unless otherwise specified, a 300 nM concentration for Osi and Lor, or a 200 ng ml^−1^ for Dox, and a 72 h treatment time were used for *in vitro* cell culture experiments.

### Doxycycline-inducible overexpression of A3A and A3B

Cells were seeded into six-well plates at a density of 2 × 10^5^ per well. After 24 hours, they were transduced with wild-type (A3A^WT^, A3B^WT^) or catalytically inactive mutant (A3A^E72A^) viral particles via spinfection at 2000 rpm for 90 minutes at 37 °C with 8 μg ml-1 polybrene (Millipore, #TR-1003-G). Following transduction, cells were incubated for 24 hours at 37°C. Cells were then selected with 600 μg/mL G418 (Gibco, #10131035) for 7 days, followed by incubation with 200 ng/mL doxycycline (Sigma) for 72 hours. Expression of Flag-A3A was confirmed by immunoblotting.

### ApoFLARE and LgBiT cells generation

#### Transient transfection

HeLa cells were transiently transfected using the following protocol: cells were plated at a density of 10^5^ cells/ml in 75 µl of DMEM + 10% FBS in two 96-well assay plates (Corning, #3917). The reporter constructs were transfected using FuGENE® 4K Transfection Reagent (Promega, #E5911), either with or without 10-fold dilution of the reporter DNA into Transfection Carrier DNA (Promega, #E4881) to yield either 50ng or 5ng reporter per well in 5µl transfection mixture, and incubated for 24 hours at 37°C. 293T, TetA3A^WT,^ and TetA3A^E72A^ PC9 were transiently transfected using the following protocol: cells were plated at a density of 2000 cells/ well in 150 µl of DMEM (293T) or RPMI (PC9) + 10% FBS in 96-well assay plates (Corning, #353377). The day after, ApoFLARE reporter constructs were transfected using Lipofectamine 3000 Transfection Reagent (Thermo Fisher, #L3000015) according to the manufacturer’s protocol and incubated for 24 hours at 37°C.

#### Stable transduction

To generate stably expressing ApoFLARE cells, the ApoFLARE reporter sequence was cloned into the pLV-EF1a-IRES-Puro lentiviral backbone using the NEBuilder® HiFi DNA Assembly Master Mix (New England Biolabs, #E2621L). Cells were seeded into 12-well plates at a density of 1 × 10^5^ per well; 24 h later they were infected with ApoFLARE reporter constructs (Reporter A substrate, Reporter A product, Reporter B substrate, Reporter B product) viral particles by spinfection at 2000 rpm for 90 min at 37 °C with 8 μg ml^−1^ polybrene (Millipore, #TR-1003-G), followed by incubation for 24 h at 37 °C. Cells were selected in 1 μg ml^−1^ Puromycin (Gibco, #A1113802) for 7 days. To generate stably expressing ApoFLARE: LgBiT cells, the LgBiT sequence was moved from the LgBiT Expression Vector (Promega, #N2681) into the pLV-EF1a-IRES-Blast lentiviral backbone using the NEBuilder® HiFi DNA Assembly Master Mix (New England Biolabs, #E2621L). ApoFLARE reporter cells were seeded into 12-well plates at a density of 1 × 10^5^ per well; 24 h later, they were infected with LgBiT viral particles by spinfection at 2000 rpm for 90 min at 37 °C with 8 μg ml^−1^ polybrene (Millipore, #TR-1003-G), followed by incubation for 24 h at 37 °C. Cells were selected in 5 μg ml^−1^ Blasticidin (Gibco, # A1113903) for 7 days.

### NanoBiT® luminescence measurement

#### Lytic assay

ApoFLARE-expressing cells were plated with either 1000 (controls) or 10000 (treatment) cells per well in appropriate media into 96-well assay plates (Corning, #353377) and incubated overnight at 37°C. The next day, cells were treated with the indicated drugs and incubated for the indicated timepoints at 37°C. The NanoBiT® and firefly luciferase signals were measured in each well using the Nano-Glo® HiBiT Dual-Luciferase® Reporter System (Promega, #NE2010) according to the manufacturer’s protocol. Briefly, ONE-Glo™ EX+LgBiT Reagent (LgBiT Protein diluted 1:100 into reconstituted ONE-Glo™ EX Reagent) was added to each well in a volume equal to the culture medium, incubated for 10 minutes, and firefly luminescence was measured. NanoDLR™ Stop & Glo® Reagent was then added in an equal volume, incubated for 10 minutes, and NanoBiT® luminescence was measured. After background subtraction, NanoBiT® signal was normalized to firefly signal for each well to calculate editing percentage as described in the Results section. Luminescence was measured using an Envision 2104 Multilabel Reader (PerkinElmer) with the Ultra-Sensitive Luminescence protocol to minimize crosstalk between wells.

#### Real-time assay

For real-time measurements in ApoFLARE: LgBiT-expressing living cells, luminescence was measured daily up to 72 hours, starting the day after seeding, using Luciferin (Revvity, #122799A) to measure Firefly signal and the Nano-Glo® Endurazine™ Live Cell Substrate (Promega, #N2570) to measure the NanoBiT® signal. Each condition was plated in duplicate to allow for measurement of both luciferases. Briefly, Endurazine™ was diluted 1:500 in medium, added to each well, and incubated for 3 hours at 37°C to reach a stable signal. Luciferin was diluted 1:100 in medium, added to each well, and incubated for 10 minutes at 37°C to reach a stable signal. Luminescence was measured using an Envision 2104 Multilabel Reader (PerkinElmer) with the Ultra-Sensitive Luminescence protocol to minimize crosstalk between wells. After background subtraction, the HiBiT signal was normalized to the firefly signal for each well, and the ApoFLARE editing % was calculated according to the formula illustrated in the Results section for each timepoint. Fresh osimertinib-containing medium was used to replace luciferin-containing wells immediately after each assay.

### RT-qPCR assay for gene expression

Cells were seeded 24 h before the addition of the drug and grown to a confluency of 80%. Cells were then treated with drugs for the indicated time points, and RNA was extracted using the RNeasy Plus Mini Kit (Qiagen, #74136). Complementary DNA was prepared from 100-1000 ng of total RNA with the High-Capacity cDNA Reverse Transcription Kit (Applied Biosystems, #4368813) using random primers. qPCR was performed using SYBR™ Select Master Mix (Applied Biosystems, #4472920) on a LightCycler 480 with analysis software v.1.5 (Roche). mRNA expression relative to control gene TBP (TATA-box binding protein) mRNA levels was calculated using the delta-delta threshold cycle (ΔΔCT) method. Primer sequences are provided in Supplementary Table 3.

### Cas9-sgRNA complex preparation

Gene knockout was performed using recombinant Cas9 ribonucleoprotein (RNP) complexes delivered via electroporation. CRISPR RNAs (crRNAs) targeting A3A and A3B are listed in Supplementary Table 2 and were synthesized by IDT. crRNAs were first incubated with 488 or 647 ATTO-labelled trans-activating crRNA (tracrRNA - IDT, #10007810 and #10007853), followed by incubation with recombinant Cas9 enzyme (IDT, #1081058) at a molar ratio of 1:1 in Neon™ NxT electroporation buffer R (Invitrogen, #N1025B) for 5 minutes at room temperature to allow complex formation. Cells were harvested at 70-80% confluence and resuspended in Neon™ NxT electroporation buffer R at a density of 1 × 10^6^ cells per reaction. Pre-assembled RNP complexes were added directly to the cell suspension and electroporated using the Neon™ NxT Electroporation System (Program A549, optimized). Immediately after electroporation, cells were transferred into pre-warmed complete RPMI medium without antibiotics and allowed to recover for 24 hours. After recovery, ATTO488-labelled cells were sorted into single-cell clones by flow cytometry to select for Cas9-RNP-containing cells. Post-recovery, individual clones were expanded and screened by direct sequencing, immunoblotting, or DDOST ddPCR following Osi treatment. Indel frequencies were quantified using TIDE analysis.

### Immunoblotting

Cells were treated with TKI (osimertinib, 1 μM for 72 hours) or doxycycline (200 ng/mL for 72 hours), followed by lysis in RIPA buffer supplemented with protease and phosphatase inhibitors. Immunoblotting was conducted as previously documented (28). The primary and secondary antibodies utilized are listed in Supplementary Table 3. Imaging of the immunoblots was performed using the G:BOX Chemi-XRQ system with GeneSys v.1.6.5.0 (Syngene) analysis software.

### Digital PCR assay

ddPCR was performed as previously reported (8). Briefly, DDOST or HiBiT Reporter A primers, designed using the Custom Assay Design Builder tool (Bio-Rad), were added along with 5-20 ng of cDNA and 2 μl of primers to PCR reactions containing ddPCR Supermix for Probes (no dUTP) in a total volume of 22 μl. Droplets were generated with a QX200 Droplet Generator (Bio-Rad). After droplet formation, the cDNA template was amplified in a C1000 Touch Thermal Cycler (Bio-Rad) under the following conditions: 5 min at 95 °C, 40 cycles of 94 °C for 30 s, 53 °C for 1 min, and a final step at 98 °C for 10 min (ramp rate of 2 °C s−1). The droplets were then analyzed with the QX200 Droplet Reader (Bio-Rad) for fluorescence detection of FAM and HEX probes, using Quanta Soft analysis software (Bio-Rad) to determine the fractional abundances of edited RNAs.

### Data and statistical analysis

Data were analyzed with GraphPad Prism software v.11.0.0 (GraphPad Software). Pairwise group comparisons, such as experimental versus control, were performed using paired or unpaired Student’s t-tests, Mann-Whitney tests, or ANOVA, as appropriate.

## RESULTS

### Engineering a dual-luciferase HiBiT-based reporter to detect APOBEC3A editing

Motivated by prior studies demonstrating that A3A efficiently deaminates specific mRNA hairpin substrates (16, 17, 20), we reasoned that embedding a preferred A3A hairpin substrate within the HiBiT-encoding RNA sequence (21) could couple A3A mRNA editing to changes in HiBiT-LgBiT complementation, forming the basis for a luminescence-based A3A editing reporter. Therefore, we sought to re-engineer the HiBiT-encoding DNA sequence with a single-nucleotide T>C substitution at the second position of a TpT dinucleotide that 1) results in an amino acid change that disrupts complementation with LgBiT and 2) is positioned within an optimal A3A substrate hairpin loop. In the presence of A3A, C>U editing at the corresponding UpC dinucleotide embedded within the mRNA hairpin loop will convert the “substrate” into the “product*”* configuration, generating a HiBiT peptide sequence that can effectively complement with the LgBiT subunit to form the active NanoBiT® luciferase (Fig. 1A). To engineer A3A-editable sites within the HiBiT coding sequence, we first identified TpT dinucleotide motifs (or UpU in the mRNA sequence) or dinucleotides for which conversion to TpT creates a synonymous substitution that does not alter the amino acid sequence (Sup. Fig. 1A). Next, we examined whether the corresponding TpC of each TpT created a non-synonymous substitution that altered the amino acid sequence. Of the eight candidate TpT motifs (Sup. Fig. 1A; Step 1, designated a-h), five encoded non-synonymous substitutions when converted to TpC (Sup. Fig. 1A; Step 2). These included amino acid changes with significant structural differences (for example, W>R, S>P), suggesting the potential for disrupting HiBiT/LgBiT complementation. For each candidate site, we optimized the surrounding sequence context by introducing synonymous substitutions to generate a hairpin consisting of a four-nucleotide loop and a six-base-pair stem while preserving the correct amino acid sequence following editing (Sup. Fig. 1A; Step 3). Three candidate sites met both structural and coding design constraints (#1-3), with one located entirely within the original HiBiT coding sequence and the other two created by adding sequence before or after the HiBiT sequence. In addition, we identified three additional TpC hairpins created by altering additional amino acids (#4-6). Finally, we introduced additional synonymous substitutions to increase stem length (7-16 bp), creating 6 potential editing reporters (Sup. Fig. 1A; Step 4, Sup. Fig. 1B).

**Figure 1.**
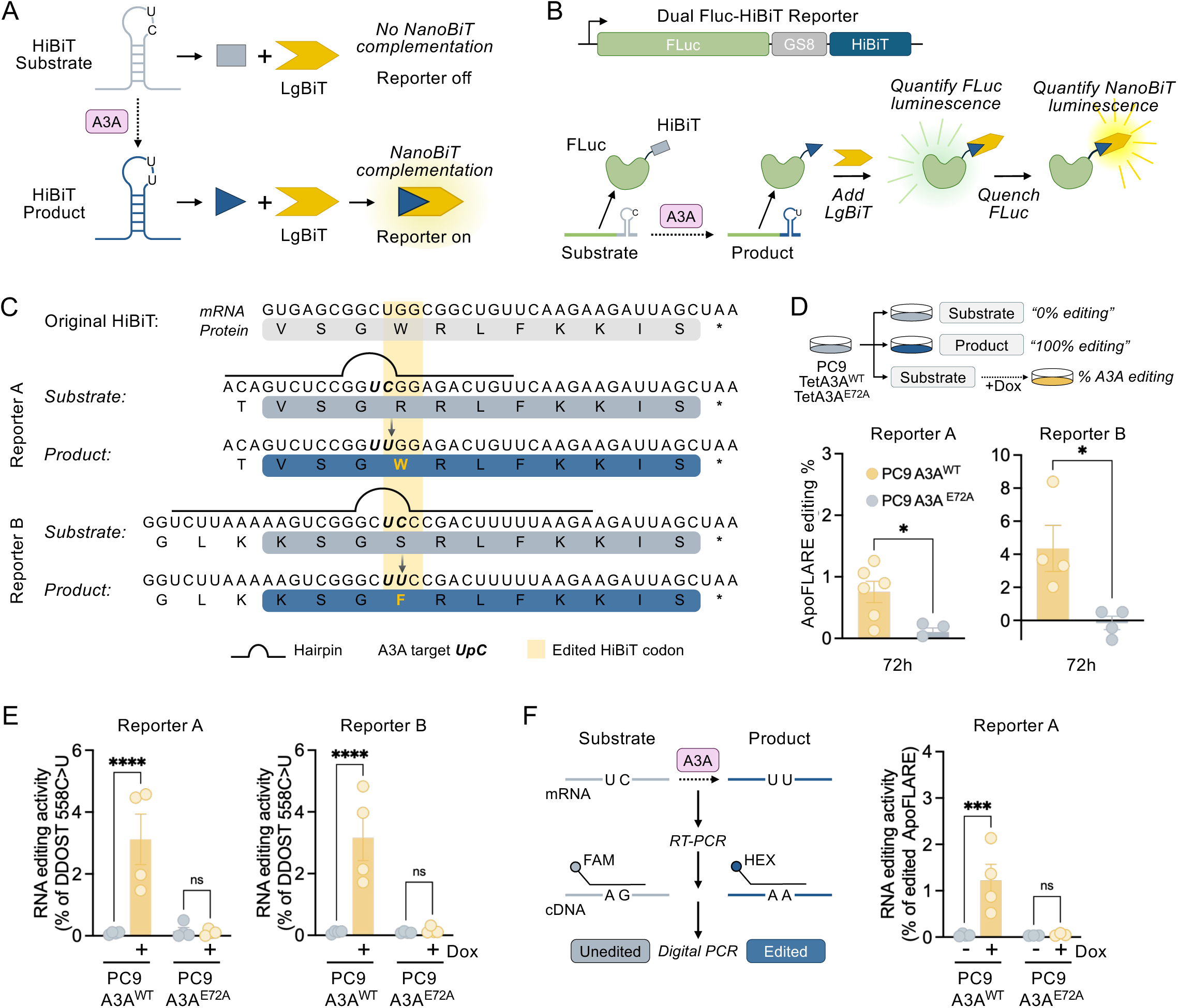
**A**. Conceptual schema of ApoFLARE reporter. A hairpin loop substrate in the HiBiT sequence can be edited by A3A to restore complementation with LgBiT and subsequent NanoBiT® luminescence. **B**. Schematic of the bicistronic reporter construct and ApoFLARE dual luminescence endpoint assay. The modified HiBiT tag sequence is fused to the C-terminal end of firefly luciferase (FLuc) with a separating linker region (GS8). FLuc luminescence is measured first, quenched, followed by measurement of NanoBiT® luminescence. **C**. Schematic showing the original HiBiT sequence (mRNA and protein) and the modified substrate (gray) and product (blue) sequences for Reporter A and Reporter B. The A3A target UpC motif is in bold, and the resulting edited HiBiT codon is highlighted in yellow. **D**. ApoFLARE Reporter A or B was transfected into PC9 TetA3A^WT^ and TetA3A^E72A^ cells; A3A editing % was determined following 72 hours of doxycycline (200 ng/mL) treatment (mean ± SEM. of four biological replicates; Student’s t-test). **E**. DDOST 558 C>U mRNA editing determined by ddPCR. (mean ± SEM of three to four biological replicates, two-way ANOVA followed by Sidak’s post-hoc test). **F**. Left, Schematic of allele-specific HiBiT ddPCR assay. Right, Reporter A mRNA editing % as determined by ddPCR. (mean ± SEM of three to four biological replicates, two-way ANOVA followed by Sidak’s post-hoc test). Dox, doxycycline.

To create the reporter constructs, we fused the re-engineered HiBiT “substrate” sequences in-frame to the C-terminus of firefly luciferase (FLuc), generating a dual-luciferase reporter expressed as a single polypeptide (Fig. 1B). This 1:1 stoichiometric configuration enables FLuc luminescence to serve as a control for variability in reporter expression. Additionally, we generated reporter constructs incorporating each of the corresponding “product” sequences (Sup. Fig. 1B). As an endpoint assay, FLuc luminescence is measured first and then quenched, followed by quantification of the HiBiT signal (generated by HiBiT/LgBiT complementation to form NanoBiT® luciferase) (Fig. 1B). HiBiT luminescence values are then normalized to FLuc activity to correct for differences in transcription, translation, and reporter abundance. The resulting HiBiT/FLuc ratios for the paired substrate and product configurations are compared to determine the percent conversion, representing the dynamic range of editing from 0% (substrate) to 100% (product), respectively.

Substrate and product reporter constructs were initially tested in HeLa cells after transient transfection with either 50 ng or 5 ng of reporter DNA per well (Sup. Fig 2A). 24 hours after transfection, we quantified firefly and HiBiT luminescence using the Nano-Glo® HiBiT Dual-Luciferase® Reporter System (HiBiT NanoDLR™) (Sup. Fig 2B, C). After background subtraction, we normalized NanoBiT® luminescence to the FLuc signal and calculated the relative fold change between the substrate and product configurations for each reporter (Sup. Fig 2D, E). With the exception of reporter #3, which showed NanoBiT®/FLuc similar to the original HiBiT, all substrate configurations showed a decreased NanoBiT®/FLuc ratio (#4, #5 < #6 < #2, #1), indicating decreased LgBiT complementation. By comparison, the product configuration corresponding to each of these five modified HiBiT sequences yielded increased NanoBiT® /FLuc ratios, ranging from 30-300 fold difference between substrate and product (#4, #2, #1 > #6, #5). Across reporters, the relative separation between substrate and product configuration was largely preserved at both DNA input levels, indicating that reporter discrimination was not strongly dependent on plasmid dosage. However, transfection with 50 ng consistently yielded higher absolute luminescence and a more robust signal-to-background ratio. We therefore used 50 ng DNA for subsequent experiments to maximize signal stability while maintaining clear configuration-dependent discrimination. To determine whether signal intensity and substrate-product discrimination were reproducible across cellular contexts, we similarly evaluated candidate reporters (#1, #2, #4, #5, #6) in 293T cells (Sup. Fig. 2F). Consistent with HeLa cells, all reporters exhibited a clear difference in NanoBiT®/FLuc ratio between substrate and product configurations (Sup. Fig 2G). Collectively, these results demonstrate that single-amino acid modifications within the HiBiT tag reduce complementation with LgBiT, thereby discriminating between “off” and “on” states that differ by a single-nucleotide substitution corresponding to a potential A3A editing motif.

We selected two ApoFLARE reporter variants for further characterization: #2 (hereafter designated Reporter A) and #6 (Reporter B). Both reporters are derived from modifications of the same codon within the HiBiT sequence but differ in the specific nucleotide edited and consequently in the encoded amino acid sequence (Fig. 1C). Reporter A encodes a TpC (substrate) > TpT (product) that results in codon 4R>W, and incorporates modifications in synonymous nucleotide positions +3, 4, 5, 9, 10, 13, 15 and 3 additional 5’ nucleotides (adding a N-terminal Thr residue) that create a 9-bp stem. Reporter B encodes a TpC (substrate) > TpT (product) that results in codon 4S>F, and also incorporates additional nucleotide modifications designed to strengthen and extend the hairpin stem (nucleotide positions +1, 2, 3 that encode for a lysine instead of a valine, synonymous nucleotide positions +4, 5, 11, 12, and 9 additional 5’ nucleotides) (Fig. 1C). To determine whether A3A-mediated editing can covert ApoFLARE reporter substrate to product and generate luminescence, we introduced Reporter A and Reporter B into PC9 cells, an *EGFR*-mutant non-small cell lung cancer (NSCLC) model with low-to-undetectable basal A3A expression (8), expressing a doxycycline-inducible wild-type A3A (PC9 TetA3A^WT^)or a catalytically-inactive A3A^E72A^ mutant (PC9 TetA3A^E72A^) (Sup. Fig. 3A). Reporter constructs were delivered by lentiviral transduction under the EF1a promoter to ensure stable expression. Following A3A induction with doxycycline, reporter activity was quantified using the HiBiT NanoDLR™ assay (Fig. 1D). We calculated the fractional conversion of the NanoBiT®/FLuc luminescence ratio from the unedited substrate configuration to the fully edited product configuration using the formula 100×(R_sample_−R_substrate_)/(R_product_−R_substrate_), where R denotes the normalized NanoBiT®/FLuc luminescence ratio (Sup. Fig. 3B). Both reporters exhibited a significant increase in NanoBiT® luminescence in the presence of wild-type A3A but not A3A^E72A^, indicating that reporter activation reflects A3A catalytic activity (Fig. 1D, Sup. Fig. 3C). Comparison of the two reporters revealed distinct performance profiles, with Reporter A displaying a broader dynamic range (~2.5 log difference between substrate and product NanoBiT®/FLuc ratio) and editing levels around 1%, while Reporter B demonstrated increased levels of A3A editing (~10%) but narrower dynamic range (~1.5 log) (Fig. 1D, Sup. Fig. 3C).

To validate that the ApoFLARE reporter reflects A3A mRNA editing, we first performed allele-specific ddPCR to quantify editing of the DDOST mRNA hotspot, a well-established assay of A3A editing activity (15, 19). Treatment with doxycycline led to DDOST editing in PC9 A3A^WT^ cells expressing Reporter A and Reporter B, but not in A3A^E72A^ cells, confirming the specificity of our reporter system in detecting A3A-mediated editing signals (Fig. 1E). Next, we developed an allele-specific ddPCR assay to directly quantify editing of Reporter A HiBiT tag transcripts (Fig. 1F, Sup. Fig. 3D) and confirmed the ability to quantify mixtures of substrate and product transcripts across a wide range of ratios (Sup. Fig. 3E). We applied the HiBiT ddPCR assay to quantify edited Reporter A transcripts in PC9 A3A^WT^ or A3A^E72A^ cells, which demonstrated editing levels that closely correlated with that calculated from the luminescence measurements, confirming that ratiometric luminescence readout reflects direct editing of the engineered HiBiT sequence in a cellular context (Fig. 1F). Together, these results validate the ApoFLARE reporter as a sensitive, specific, and quantitative readout for A3A editing activity in cells.

### ApoFLARE detects endogenous A3A editing

To determine whether ApoFLARE can detect endogenous A3A editing, we assessed several biologically relevant contexts in which A3A is induced by exogenous stimuli. First, we examined THP-1 cells, a physiologic model of A3A induction in response to interferon stimulation (22, 23). We treated THP1 ApoFLARE reporter cells with interferon-α (IFNα) and measured A3A mRNA expression and reporter editing activity at 6 and 24 hours following treatment (Fig. 2A). IFNα exposure resulted in a time-dependent increase in A3A mRNA (Fig. 2B), which corresponded to a progressive increase in ApoFLARE reporter editing: ~0.5% for Reporter A and up to ~4% for Reporter B (Fig. 2C). Consistent with reporter activation, IFNα treatment also increased editing of the endogenous A3A substrate DDOST, as measured by allele-specific ddPCR (Fig. 2D). By comparison, A3B expression remained low and was not detectably induced by IFNα (Fig. 2B), consistent with A3A-dependent reporter activation.

**Figure 2.**
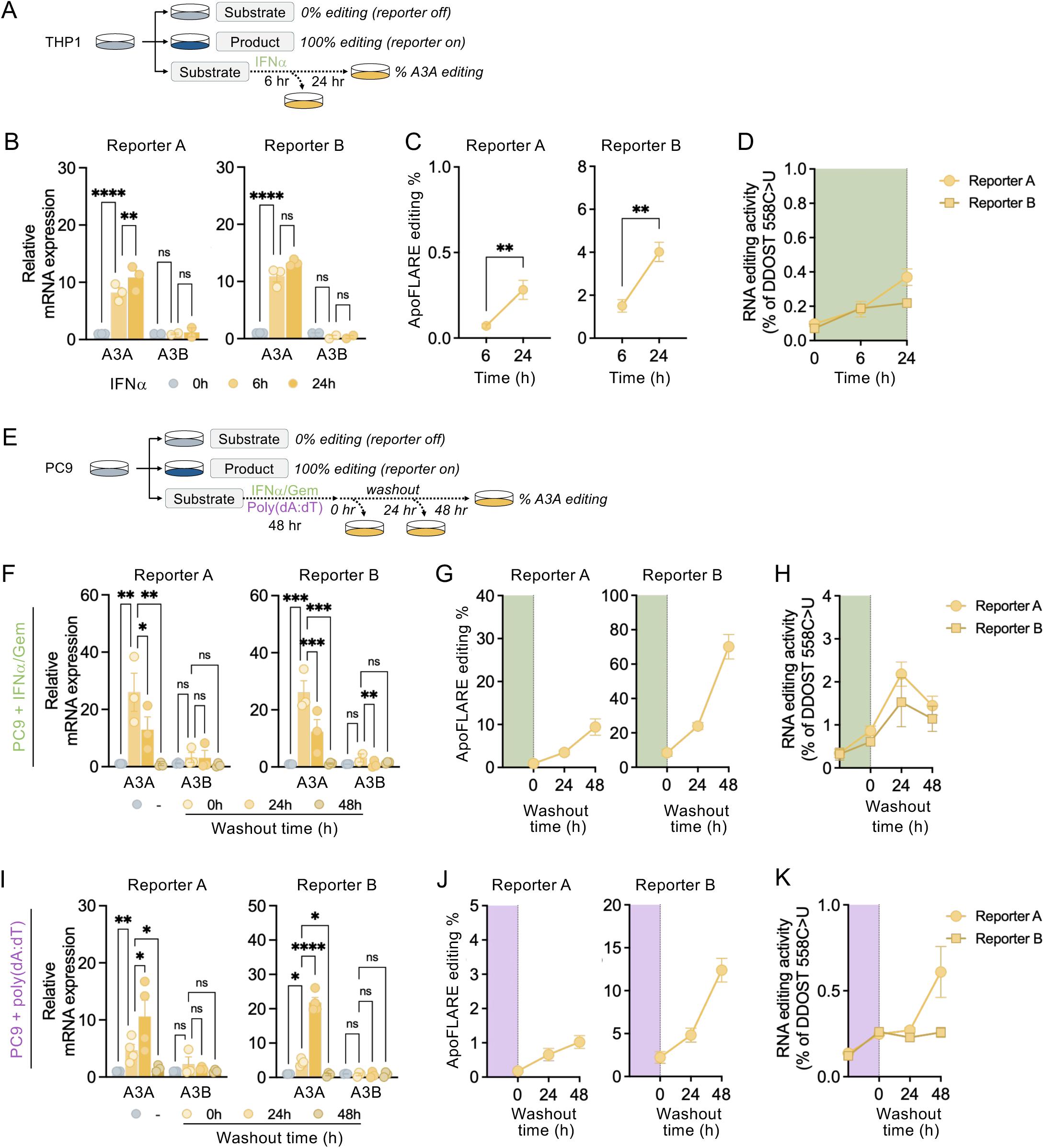
**A**. Experimental schema for ApoFLARE detection of IFNα-induced A3A editing. **B**. A3A and A3B expression induced in THP1 cells upon IFNα treatment (mean ± SEM of three biological replicates, two-way ANOVA followed by Sidak’s post-hoc test). **C**. ApoFLARE editing (Reporter A and B) following 6 or 24 hours of IFNα treatment (mean ± SEM of three biological replicates; Student’s t-test). **D**. DDOST 558 C>U mRNA editing determined by ddPCR. (mean ± SEM of three biological replicates). **E**. Experimental schema for ApoFLARE detection of IFNα/gemcitabine or poly(dA:DT)-induced A3A editing in PC9 cells. **F**. A3A and A3B expression in PC9 cells treated with IFNα/Gem and released into drug-free media (mean ± SEM of three biological replicates, two-way ANOVA followed by Sidak’s post-hoc test). **G**. ApoFLARE editing (Reporter A and B) following IFNα/Gem washout (mean ± SEM of two biological replicates). **H**. DDOST 558 C>U mRNA editing determined by ddPCR. (mean ± SEM of two biological replicates). **I**. A3A and A3B expression in PC9 cells treated with poly(dA:dT) and released into drug-free media (mean ± SEM of three biological replicates, two-way ANOVA followed by Sidak’s post-hoc test). **J**. ApoFLARE editing (Reporter A and B) following poly(dA:dT) washout (mean ± SEM of two biological replicates). **K**. DDOST 558 C>U mRNA editing determined by ddPCR. (mean ± SEM of two biological replicates). IFNα, interferon-α; Gem, gemcitabine.

Second, we exposed PC9 ApoFLARE reporter cells to gemcitabine and IFNα, stress conditions previously shown to induce A3A expression and activity (15). We treated cells for 48h and then cultured in drug-free media, assessing NanoBiT® luminescence and A3A expression at the end of treatment and at 24-hour intervals up to 48h after drug washout (Fig. 2E). A3A mRNA levels peaked after IFNα/Gem exposure, then subsided over 48 hours of drug washout (Fig. 2F). ApoFLARE reporter editing activity exhibited delayed kinetics, with a modest increase in the time of drug washout and further increase over the following 48 hours (Fig. 2G). DDOST RNA editing exhibited comparable delayed kinetics, increasing after drug washout and mirroring the temporal pattern observed with ApoFLARE reporter activation (Fig. 2H). This persistence of reporter editing despite declining A3A transcript levels is consistent with prior observations that A3A enzymatic activity and RNA editing can outlast transient A3A transcription due to greater stability of the protein relative to its mRNA (15), demonstrating that ApoFLARE can capture episodic A3A activation and sustained editing activity in tumor cells.

Third, we performed an analogous set of experiments using transfected poly(dA:dT), a synthetic double-stranded DNA that activates cytosolic DNA-sensing pathways and induces a type I interferon response, which, in turn, stimulates A3A expression. PC9 ApoFLARE reporter cells were treated with poly(dA:dT), followed by release into DNA-free media, and A3A expression and reporter activity were monitored (Fig. 2E). Poly(dA:dT) exposure induced expression of A3A mRNA levels (Fig. 2I) and ApoFLARE reporter activation (Fig. 2J). A3A transcript levels peaked 24 hours after stimulus withdrawal then declined, whereas ApoFLARE reporter editing activity continued to increase through the washout period. Endogenous DDOST editing similarly persisted following stimulus withdrawal (Fig. 2K), further demonstrating that ApoFLARE quantitatively reflects sustained A3A catalytic activity in this context. Collectively, these findings demonstrate that ApoFLARE reports endogenous A3A editing activity, including in contexts where A3A expression is transiently induced.

### Targeted therapy-induced A3A activation is detected by ApoFLARE

We previously reported that induction of A3A in lung cancer cells treated with targeted therapies accelerates the development of drug resistance (8). To determine whether ApoFLARE might enable detection of therapy-induced A3A editing in this context, we treated PC9 cells with the EGFR inhibitor osimertinib (Osi) for 72 hours (Fig. 3A). Osimertinib treatment robustly activated both Reporter A and Reporter B, as demonstrated both by luminescence readouts (Fig. 3B) and by allele-specific HiBiT ddPCR for Reporter A (Fig. 3C). Reporter activation coincided with increased A3A mRNA expression in the treated substrate (Sup. Fig. 4A) and with editing of the endogenous A3A substrate DDOST (Fig. 3D). In some cell contexts, targeted therapies can also induce A3B (9), so to confirm that the ApoFLARE signal reflects A3A editing, we generated A3A^KO^, A3B^KO^, and scramble control PC9 cells by CRISPR-mediated gene knockout, established single-cell clones (A3A^KO^#1, A3A^KO^#2, A3B^KO^#1) (Fig. 3E, Sup. Fig 4B, C) and generated stable-expressing ApoFLARE reporter cell lines. As expected, osimertinib induced DDOST editing and ApoFLARE activation in parental PC9 cells, however both responses were absent in A3A^KO^ cells (Fig. 3G, H). In contrast, A3B^KO^ cells exhibited both osimertinib-induced DDOST editing and activation of Reporter A and Reporter B. (Fig. 3H, I). In addition, inducible expression of wild-type A3B in PC9 cells (PC9 TetA3B^WT^) did not produce any detectable increase in luminescent signal, reinforcing the specificity of ApoFLARE for A3A-mediated editing (Sup. Fig. 4D-F). These data demonstrate that ApoFLARE selectively detects osimertinib-induced A3A editing and not A3B activity, consistent with the distinct substrate preferences of these closely related cytidine deaminases.

**Figure 3.**
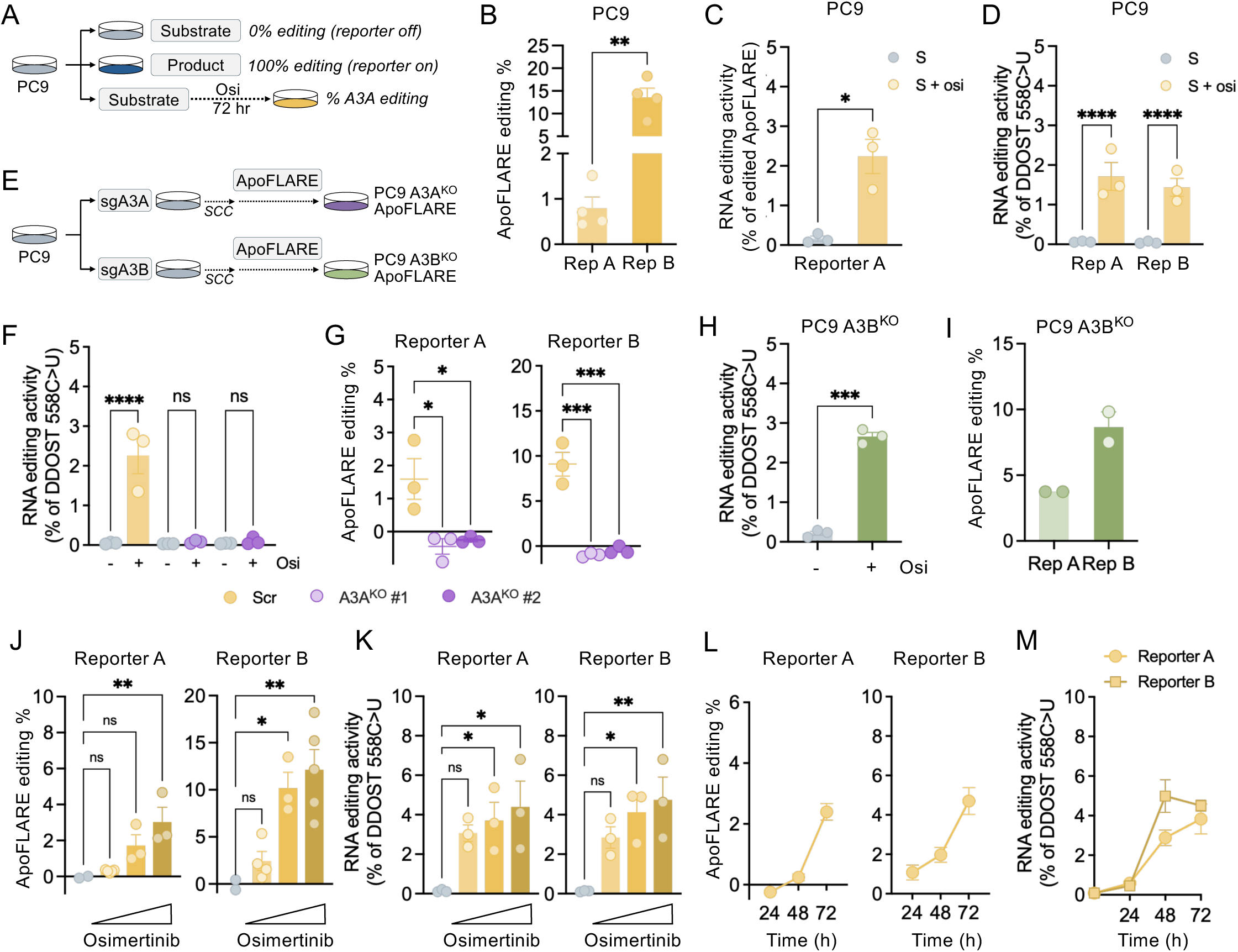
**A.** Experimental schema for ApoFLARE detection of osimertinib-induced A3A editing. **B**. ApoFLARE editing (Reporter A and B) in PC9 cells after 72 hours of osimertinib (300 nM) treatment (mean ± SEM of four biological replicates; Student’s t-test). **C**. Reporter A mRNA editing determined by ddPCR (mean ± SEM of three biological replicates, Student’s t test). **D**. DDOST 558 C>U mRNA editing determined by ddPCR. (mean ± SEM of three biological replicates, two-way ANOVA followed by Sidak’s post-hoc test). **E**. Experimental schema for generation of PC9 A3A^KO^ and A3B^KO^ ApoFLARE cell lines (SCC, single cell clone). **F**. DDOST 558 C>U mRNA editing in PC9 A3A^Scr^ and PC9 A3A^KO^ single cell clones #1 and #2 after treatment with vehicle or osimertinib as determined by ddPCR. (mean ± SEM of three biological replicates, two-way ANOVA followed by Sidak’s post-hoc test). **G**. ApoFLARE editing (Reporter A and B) after treatment with osimertinib (mean ± SEM of three biological replicates; Student’s t-test). **H**. DDOST 558 C>U mRNA editing in PC9 A3B^KO^ cells after treatment with vehicle or osimertinib as determined by ddPCR. (mean ± SEM of three biological replicates, Student’s t test). **I**. ApoFLARE editing (Reporter A and B) in PC9 A3B^KO^ cells after treatment with osimertinib (mean ± SEM of two biological replicates). **J**. ApoFLARE editing (Reporter A and B) after treatment with increasing concentrations of osimertinib (0, 10 nM, 100 nM, 1 µM) for 72 hours (mean ± SEM of three to five biological replicates, one-way ANOVA followed by Sidak’s post-hoc test). **K**. DDOST 558 C>U mRNA editing determined by ddPCR after treatment with osimertinib (mean ± SEM of three biological replicates, two-way ANOVA followed by Sidak’s post-hoc test). **L**. ApoFLARE editing (Reporter A and B) after treatment with osimertinib (300 nM) for the indicated timepoints (mean ± SEM of three to five biological replicates). **M**. DDOST 558 C>U mRNA editing determined by ddPCR after treatment with osimertinib (mean ± SEM of three biological replicates). Osi, osimertinib.

Next, we assessed the ability of ApoFLARE to report dose-dependent and temporal dynamics of therapy-induced A3A activation. Increasing osimertinib concentrations (0-1 μM) drove a graded increase in ApoFLARE reporter activation (Fig. 3J), reflecting the dose-dependent induction of A3A expression and DDOST editing activity previously described following EGFR inhibition (Fig. 3K, Sup. Fig. 3G) (8). In addition, ApoFLARE reporter editing increased progressively over time and reached maximal levels after 3 days of continuous osimertinib treatment (300 nM) (Fig. 4L), closely mirroring the kinetics of A3A transcript induction and DDOST editing (Fig. M, Sup. Fig. 4H). Importantly, allele-specific ddPCR for Reporter A closely matched the editing percentages calculated from luminescence values in both drug-titration and time-course experiments, thereby further reinforcing the robustness of the reporter (Sup. Fig 4I, 4J). Together, these data show that ApoFLARE faithfully reports therapy-induced A3A editing in PC9 cells, recapitulating previously established responses to EGFR inhibition.

**Figure 4.**
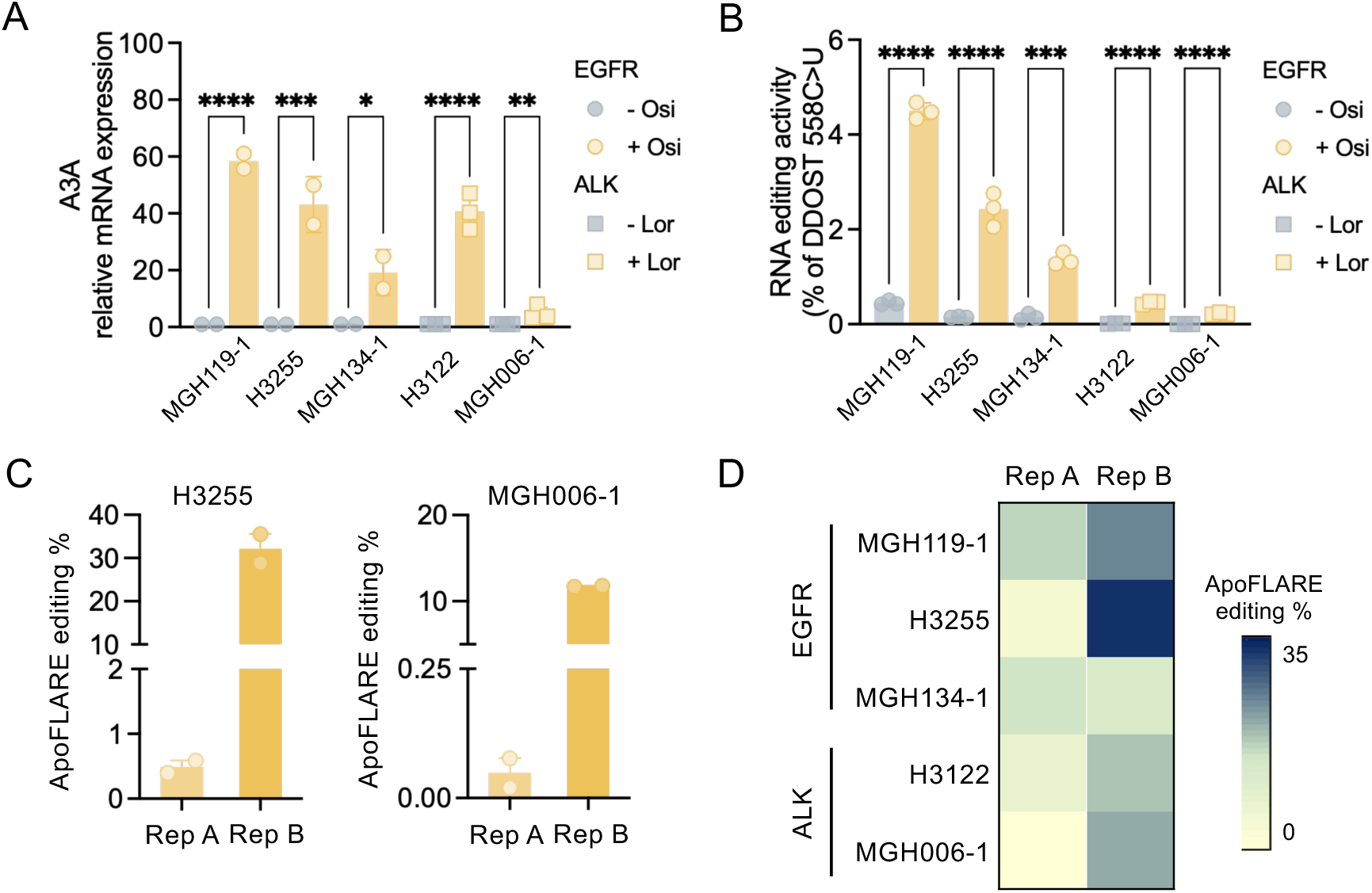
**A**. A3A expression in a panel of EGFR (MGH119-1, H3255, MGH134-1) and ALK (H3122, MGH006-1) NSCLC cell lines treated with the corresponding TKI (Osi, EGFR; Lor, ALK) for 72 hours (mean ± SEM of two to three biological replicates, two-way ANOVA followed by Sidak’s post-hoc test). **B**. DDOST 558 C>U mRNA editing determined by ddPCR. (mean ± SEM of three biological replicates, two-way ANOVA followed by Sidak’s post-hoc test). **C, D**. Bar plots and heatmap showing ApoFLARE editing in the indicated cell lines. (mean ± SEM of two to five biological replicates). TKI, tyrosine kinase inhibitor; Osi, osimertinib; Lor, lorlatinib.

To test ApoFLARE performance across a heterogeneous NSCLC context, we selected a panel of EGFR- and ALK-driven cell lines that represent variability of A3A expression observed across patient tumors (Fig. 4A). Endogenous A3A editing of the DDOST hotspot generally correlated with A3A transcript induction following the appropriate targeted therapy (osimertinib for *EGFR*-mutant models and lorlatinib (Lor) for *ALK*-rearranged models) (Fig. 4B). To determine whether ApoFLARE could report A3A activity across this range of activation states, including lower levels of A3A induction, we generated stable reporter cell lines for each model and measured luminescence under basal conditions and following treatment with corresponding tyrosine kinase inhibitors (TKIs). As expected, ApoFLARE detected robust therapy-induced reporter editing in all models tested, including those with relatively high A3A expression and DDOST editing (such as H3255), but also those with relatively low levels of A3A induction and DDOST editing (such as MGH006-1) (Fig. 4C, D). These results indicate that ApoFLARE is a sensitive reporter of A3A activity across heterogeneous oncogenic backgrounds and a wide range of activation states.

### ApoFLARE enables quantitative real-time monitoring of A3A editing in living cells

Next, we sought to extend ApoFLARE to live-cell applications and longitudinal monitoring of A3A activity in the same cell population over time. We generated PC9 reporter cell lines with stable co-expression of ApoFLARE and LgBiT, enabling intracellular NanoBiT® complementation without exogenous LgBiT supplementation. Because editing calculations require normalization of the NanoBiT® signal to Firefly luciferase to account for variability in ApoFLARE expression and cell number (which can be affected by drug treatment), we implemented a parallel dual-luciferase approach. In one set, Nano-Glo® Endurazine™ Live Cell Substrate was added at the time of osimertinib treatment to continuously monitor NanoBiT® luminescence. In the parallel set, D-luciferin, the substrate for FLuc, was added at each timepoint to quantify FLuc luminescence independently of NanoBiT® luciferase (Fig. 5A). After luminescence acquisition, luciferin-containing medium was replaced with fresh osimertinib-containing media, while Endurazine™-containing wells remained undisturbed. We calculated ApoFLARE editing at each timepoint using the same ratiometric framework established for endpoint lytic assays (Sup. Fig 3B). Using this configuration, we observed a time-dependent increase in NanoBiT® luminescence following osimertinib treatment, with ApoFLARE editing rising progressively over the monitoring period to a peak at 72 hours (Fig. 5B). Parallel RNA, collected from identically treated wells, demonstrated a concordant increase in DDOST RNA editing over time, confirming that the live-cell luminescence reflects endogenous A3A activity (Fig. 5C) Importantly, the calculated percentage of ApoFLARE editing in live cells closely matched the editing levels obtained from corresponding lytic dual-luciferase time course assays (Fig. 3L), indicating that the live-cell configuration maintains comparable detection sensitivity and quantitative performance. Therefore, ApoFLARE enables longitudinal, non-destructive quantification of A3A editing dynamics in live cells.

**Figure 5.**
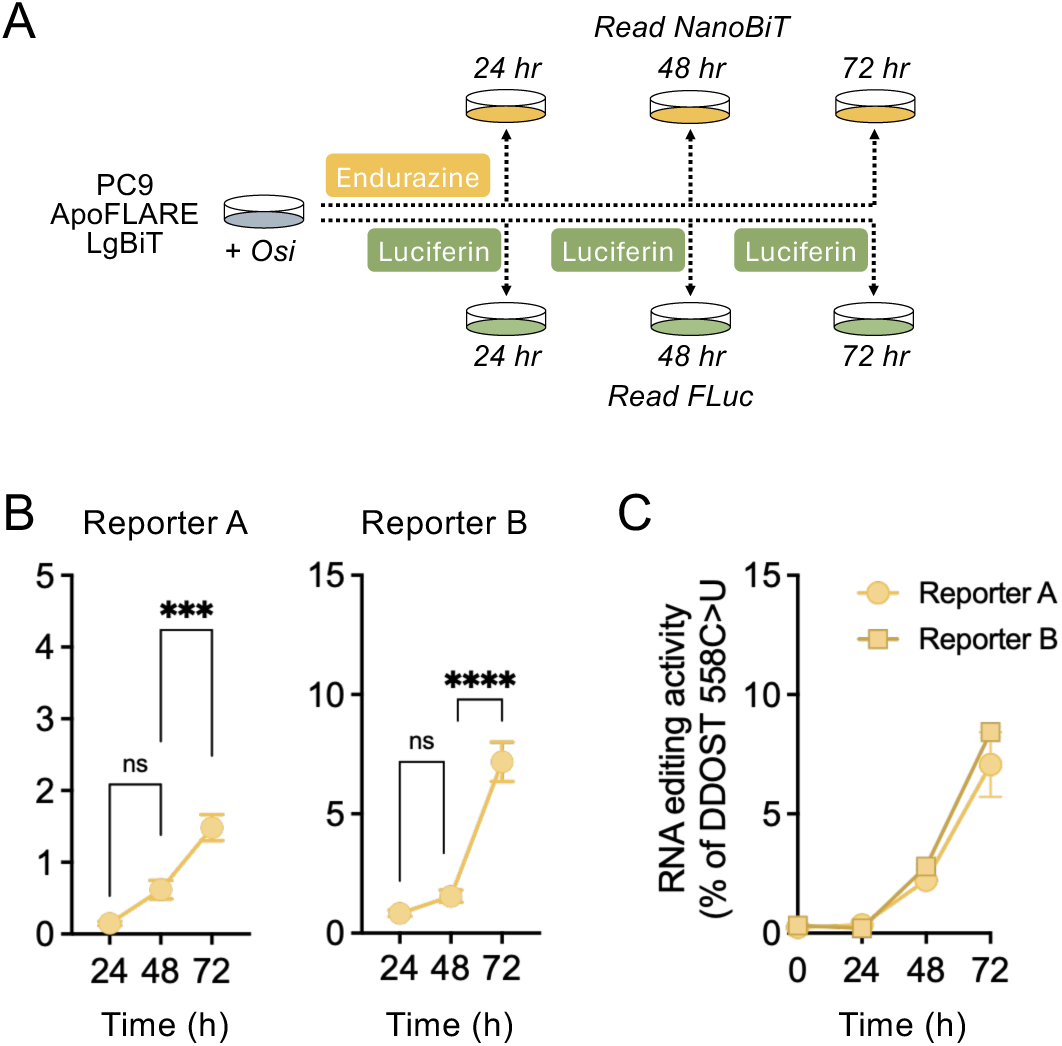
**A**. Experimental schema for dual-luciferase live-cell imaging. **B**. ApoFLARE editing (Reporter A and B) in PC9 cells treated with osimertinib (300 nM) for the indicated timepoints (mean ± SEM of three biological replicates, two-way ANOVA followed by Sidak’s post-hoc test). **C**. DDOST 558 C>U mRNA editing in PC9 cells treated with osimertinib determined by ddPCR. (mean ± SEM of three biological replicates). Osi, osimertinib.

## DISCUSSION

APOBEC3A is increasingly recognized as a dynamic and stress-responsive driver of mutagenesis in cancer (5, 6, 13, 14). However, tools capable of directly measuring cellular editing activity in real time have remained limited. Here, we describe ApoFLARE, a genetically-encoded reporter that translates A3A-mediated cytidine deamination into a quantifiable luminescent signal, allowing direct assessment of enzymatic activity rather than relying on expression levels or retrospective mutational signatures. ApoFLARE leverages a defining biochemical feature of A3A, its strong preference for structured single-stranded hairpin substrates, to couple site-specific RNA editing to NanoBiT® complementation. Reporter activation required catalytic A3A activity and was abolished in A3A-deficient cells, but remained intact in A3B-knockout backgrounds, thereby confirming specificity for A3A over A3B. This distinction is particularly relevant given the overlapping sequence similarity yet divergent regulation of APOBEC3A and APOBEC3B and offers the capability to selectively resolve A3A-mediated editing in tumor systems.

Orthogonal validation showed that luminescent output quantitatively tracks endogenous editing under stress and therapeutic conditions, including DDOST RNA editing, the current gold-standard assay for A3A editing activity (15). Notably, ApoFLARE detected sustained editing even as A3A transcript levels decreased. This finding supports previous hotspot-based RNA editing studies indicating that A3A catalytic output can continue beyond mRNA induction and that transcript abundance alone is not a reliable predictor of ongoing deamination activity (15). Therefore, the reporter reflects the functional catalytic output rather than merely indicating mRNA levels. ddPCR-based mRNA hotspot quantification is a highly sensitive and accurate method for measuring A3A-mediated RNA editing, however the multi-step workflow limits scalability and temporal resolution in large-scale or long-term experiments. In contrast, ApoFLARE can be read via plate reader-based assay that allows rapid, parallel processing of many samples with minimal handling. While ddPCR provides nucleotide-level quantification, ApoFLARE complements this by enabling higher throughput and time-resolved functional measurement of A3A activity.

Applying ApoFLARE to oncogene-driven lung cancer models detected therapy-induced A3A editing across heterogeneous cellular contexts and revealed inter-model variability in editing magnitude. In oncogene-driven NSCLC, targeted therapies have been shown to trigger a broad, yet heterogeneous A3A activation program within drug-tolerant persister cells, promoting evolutionary diversification under therapy (8). Genetic knockout confirmed that reporter activation caused by targeted therapy is driven solely by A3A. These results demonstrate that ApoFLARE accurately reports known biological behaviors and offers a scalable, quantitative platform for perturbation studies.

A key feature of ApoFLARE is its compatibility with long-term live-cell measurements. In its current configuration, ApoFLARE allows for continuous tracking of editing kinetics at the population level. With further optimization, ApoFLARE could be adapted for live-cell bioluminescent imaging workflows, enabling spatial and temporal analysis of A3A activity in intact cells. Achieving reliable single-cell detection will likely require further refinement of reporter configuration, signal amplification, and substrate delivery to improve dynamic range and signal-to-noise. Live-cell imaging might allow direct visualization of heterogeneous or pulsatile A3A activation within specific subpopulations under therapeutic or stress conditions. Such approaches could clarify whether editing activity is uniformly induced or limited to discrete cellular states that drive adaptive evolution and clonal diversity.

Beyond mechanistic interrogation, ApoFLARE offers a basis for translational research. The system may facilitate systematic screening for A3A inhibitors, identification of cellular contexts suitable for A3A editing activation, and enable real-time monitoring of therapy-induced deamination dynamics. By enabling direct, quantitative, and time-resolved measurement of therapy-related A3A activity, ApoFLARE broadens the experimental tools available for studying APOBEC-driven mutagenesis. When combined with genomic, transcriptomic, and functional methods, this platform may help develop a more comprehensive and mechanistic understanding of how episodic deamination influences tumor evolution and therapeutic response.

## ACKNOWLEDGEMENTS

We thank members of the Hata Lab and the MGH Thoracic Oncology group for helpful feedback and support. We thank Professor Rémi Buisson and Professor Reuben Harris for providing fundamental materials used in this study.

## AUTHOR CONTRIBUTIONS

*M.V.D.M.,* Investigation, Formal analysis, Methodology, Validation, Data Curation, Writing – original draft; *B.L.B.*, Investigation; *C.T.E.*, Conceptualization, Methodology, Writing – review & editing; *A.N.H.*, Conceptualization, Methodology, Supervision, Funding acquisition, Writing – review & editing.

## SUPPLEMENTARY DATA

Supplementary data are available at NAR online.

## CONFLIC OF INTEREST

*A.N.H*. has received grants/research support from Amgen, BBOT, Bristol-Myers Squibb, C4 Therapeutics, Eli Lilly, Immuto Scientific, Novartis, Nuvalent, Pfizer, Scorpion Therapeutics, Triana Biomedicines; consulting fees from Chugai, Kerna Ventures, Nuvalent, Amgen, Pfizer.

*C.T.E.* and *B.L.B.* are employees of Promega Corporation, which commercially owns and sells the NanoBiT® technology and associated detection reagents described in this work, including modifications made to the HiBiT nucleic acid sequence.

## DATA AVAILABILITY

The data underlying this article are available from the corresponding author upon request.

## Supplemental Files

- **Supplementary Figure 1**
- **Supplementary Figure 2**
- **Supplementary Figure 3**
- **Supplementary Figure 4**
- **Supplementary Table 1**
- **Supplementary Table 2**
- **Supplementary Table 3**

## Supplementary Figure Legends

**Supplementary Figure 1. A**. Schematic showing the original HiBiT sequence (DNA, mRNA and encoded protein) and the subsequent modifications to obtain ApoFLARE reporters. Step 1, Eight candidate synonymous TpT dinucleotides in the HiBit sequence (designated a-h). Step 2, TpT → TpC modifications (blue) that resulted in codon changes (yellow). Letter designations correspond to dinucleotides in Step 1. Step 3, Nucleotide changes/additions (blue) to generate 6-bp hairpin stems within the HiBiT sequence (gray shading). Step 4, Additional nucleotides to increase stem length. **B**. Final 6 ApoFLARE structures selected for further testing. Gray – *substrate*; blue – *product*.

**Supplementary Figure 2. A**. Experimental schema for testing ApoFLARE reporter constructs in HeLa cells. **B, C**. NanoBiT®-to-FLuc luminescence ratios following transfection of either 5 or 50ng of the 6 candidate ApoFLARE reporters into HeLa cells (mean ± SEM of two technical replicates). **D, E**. Fold change in normalized substrate to product luminescent signal in HeLa cells (mean ± SEM of two technical replicates) **F**. Experimental schema for testing ApoFLARE reporter constructs in 293T cells. **G**. HiBiT-to-FLuc luminescence ratios following transfection of 5 candidate ApoFLARE reporters into 293T cells (mean ± SEM of five technical replicates). **H**. Fold change in normalized substrate to product luminescent signal in 293T cells. (mean ± SEM of five technical replicates).

**Supplementary Figure 3. A**. Immunoblot of A3A expression following doxycycline induction at the indicated timepoints in PC9 TetA3A^WT^ or TetA3A^E72A^ cells. β-actin was used as a loading control. **B**. Formula used to calculate the fractional conversion of the NanoBiT® /FLuc luminescence ratio from the unedited substrate configuration to the fully edited product configuration. R, normalized NanoBiT® /FLuc luminescence ratio. **C**. NanoBiT®-to-FLuc luminescence ratios in PC9 TetA3A^WT^ treated with doxycycline for 72 hours. Light gray, detection range from R_substrate_ (0%) to R_product_ (100%). **D**. ddPCR dotplots showing fluorescent amplitude of Reporter A and B ddPCR probes using plasmid DNA. Gray, substrate; Light gray, empty droplets; Blue, product. **E**. Detection range for allele-specific HiBiT ddPCR for ApoFLARE reporter A in mixing experiments using the indicated proportions of substrate and product plasmids.

**Supplementary figure 4. A.** A3A expression in PC9 cells treated with osimertinib for 72 hours (mean ± SEM of three biological replicates, Student’s t test). **B, C**. Immunoblot of A3A (**B**) or A3B (**C**) expression following genetic knockout after CRISPR-RNP delivery. A3A^KO^ cells were treated with vehicle or osimertinib. β-actin was used as a loading control. **D**. Experimental schema for testing ApoFLARE in PC9 cells with inducible A3B. **E, F**. ApoFLARE editing (Reporter A and B) in PC9 TetA3B^WT^ cell lines treated with dox for 72 hours, and corresponding DDOST 558 C>U mRNA editing determined by ddPCR. (mean ± SEM of two biological replicates). **G, H**. A3A expression in PC9 cells treated with increasing concentrations of osimertinib (0, 10 nM, 100 nM, 1 µM) for 72 hours (**G**) or osimertinib (300 nM) for the indicated timepoints (**H**) (mean ± SEM of three to six biological replicates, one-way ANOVA followed by Sidak’s post-hoc test). **I, J**. ApoFLARE reporter A mRNA editing determined by allele-specific ddPCR. (mean ± SEM of two to three biological replicates; **I**, one-way ANOVA followed by Sidak’s post-hoc test). Dox, doxycycline; osi, osimertinib.

