## Supplementary Figures and Legends for "ApoFLARE: a luminescent reporter for direct quantification of APOBEC3A editing activity"

Supplementary Figure 1

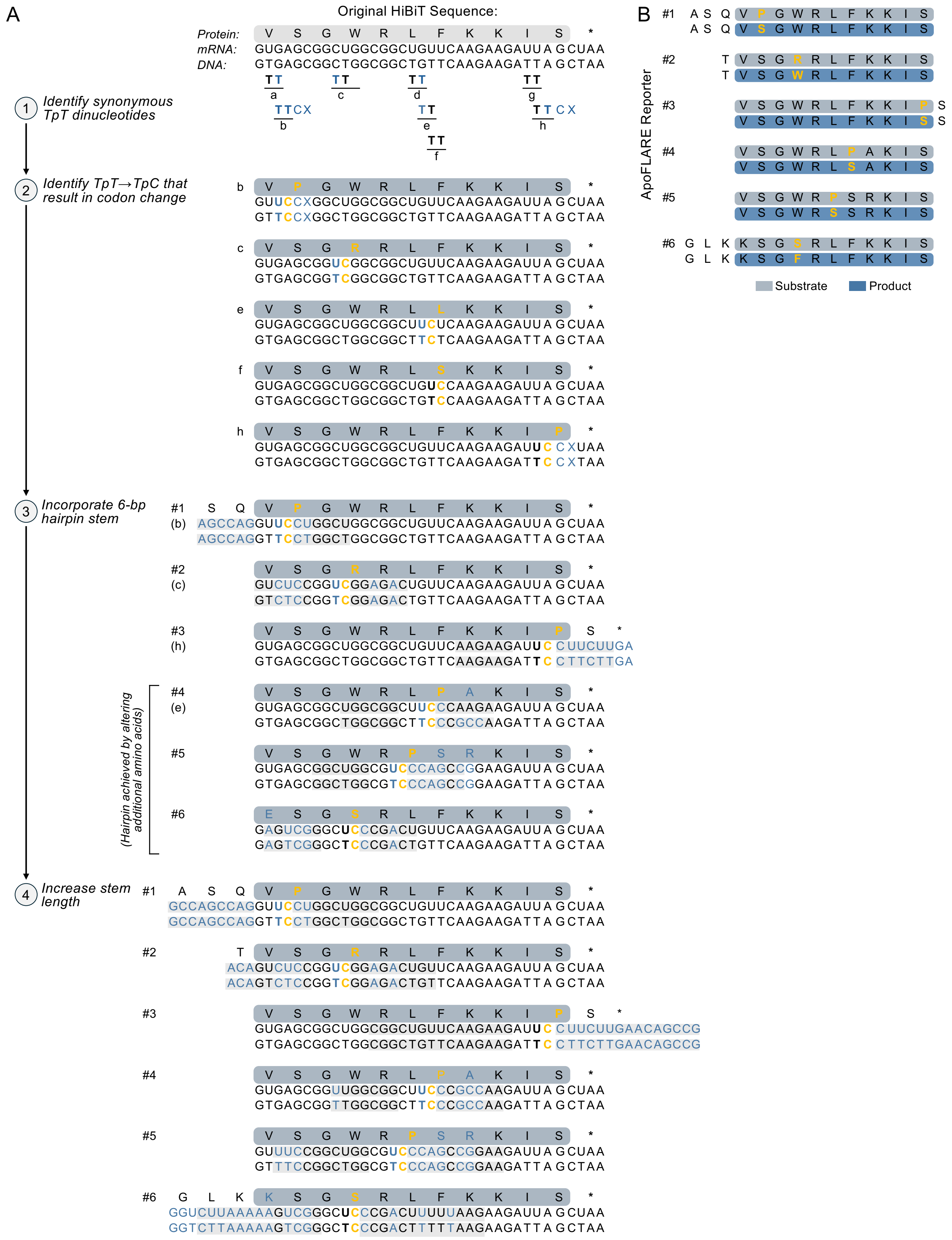

**Supplementary Figure 1. A**. Schematic showing the original HiBiT sequence (DNA, mRNA and encoded protein) and the subsequent modifications to obtain ApoFLARE reporters. Step 1, Eight candidate synonymous TpT dinucleotides in the HiBit sequence (designated a-h). Step 2, TpT → TpC modifications (blue) that resulted in codon changes (yellow). Letter designations correspond to dinucleotides in Step 1. Step 3, Nucleotide changes/additions (blue) to generate 6-bp hairpin stems within the HiBiT sequence (gray shading). Step 4, Additional nucleotides to increase stem length. **B**. Final 6 ApoFLARE structures selected for further testing. Gray – *substrate*; blue – *product*.

Supplementary Figure 2

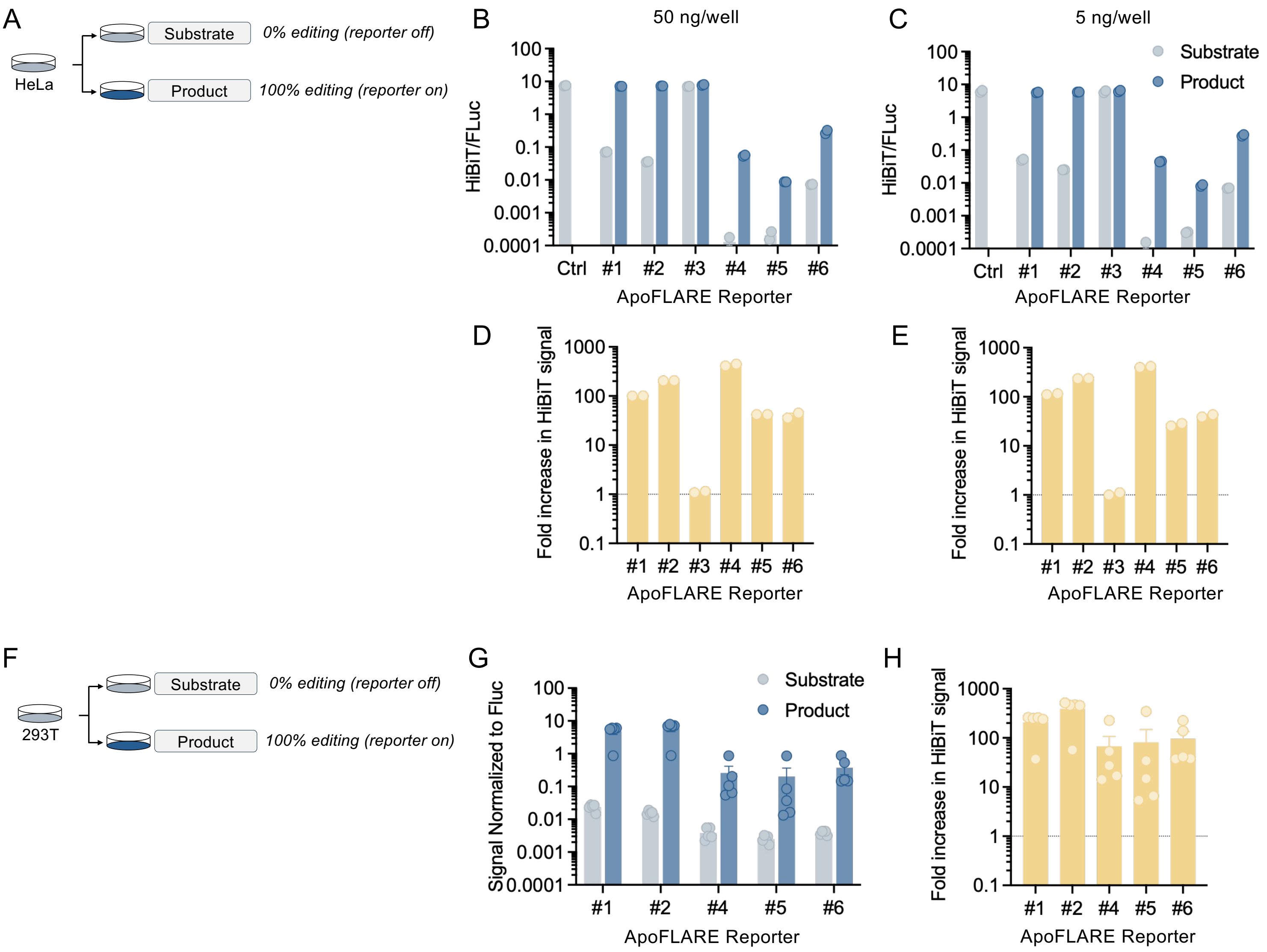


**Supplementary Figure 2. A**. Experimental schema for testing ApoFLARE reporter constructs in HeLa cells. **B, C**. NanoBiT®-to-FLuc luminescence ratios following transfection of either 5 or 50ng of the 6 candidate ApoFLARE reporters into HeLa cells (mean ± SEM of two technical replicates). **D, E**. Fold change in normalized substrate to product luminescent signal in HeLa cells (mean ± SEM of two technical replicates) **F**. Experimental schema for testing ApoFLARE reporter constructs in 293T cells. **G**. HiBiT-to-FLuc luminescence ratios following transfection of 5 candidate ApoFLARE reporters into 293T cells (mean ± SEM of five technical replicates). **H**. Fold change in normalized substrate to product luminescent signal in 293T cells. (mean ± SEM of five technical replicates).

### Supplementary Figure 3

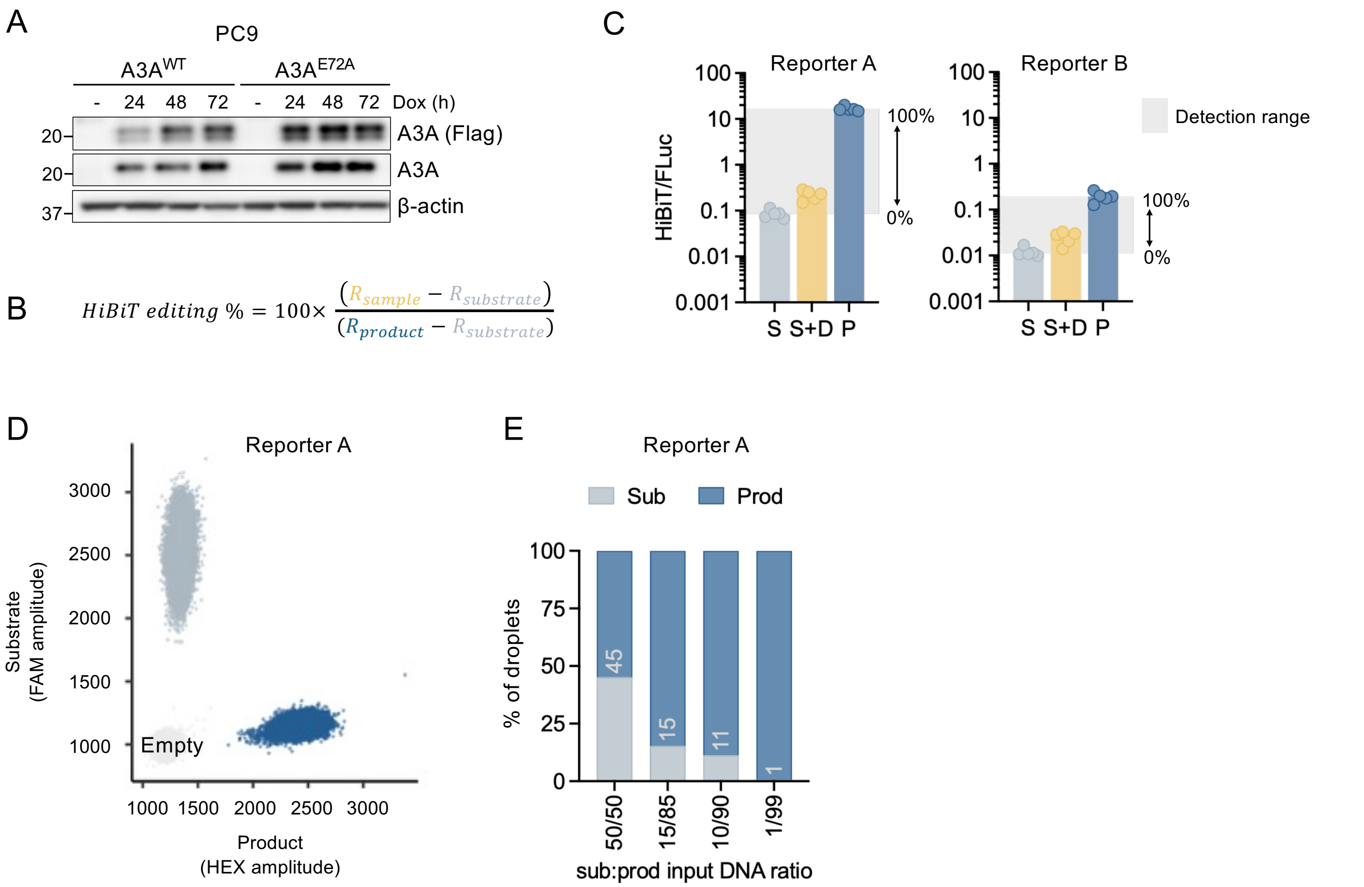


$$HiBiT\ editing\ \% = 100\times\frac{(R_{sample} - R_{substrate})}{(R_{product} - R_{substrate})}$$

**Supplementary Figure 3. A**. Immunoblot of A3A expression following doxycycline induction at the indicated timepoints in PC9 TetA3A$^{WT}$ or TetA3A$^{E72A}$ cells. β-actin was used as a loading control. **B**. Formula used to calculate the fractional conversion of the NanoBiT® /FLuc luminescence ratio from the unedited substrate configuration to the fully edited product configuration. R, normalized NanoBiT® /FLuc luminescence ratio. **C**. NanoBiT® -to-FLuc luminescence ratios in PC9 TetA3A$^{WT}$ treated with doxycycline for 72 hours. Light gray, detection range from $R_{substrate}$ (0%) to $R_{product}$ (100%). **D**. ddPCR dotplots showing fluorescent amplitude of Reporter A and B ddPCR probes using plasmid DNA. Gray, substrate; Light gray, empty droplets; Blue, product. **E**. Detection range for allele-specific HiBiT ddPCR for ApoFLARE reporter A in mixing experiments using the indicated proportions of substrate and product plasmids.

Supplementary Figure 4

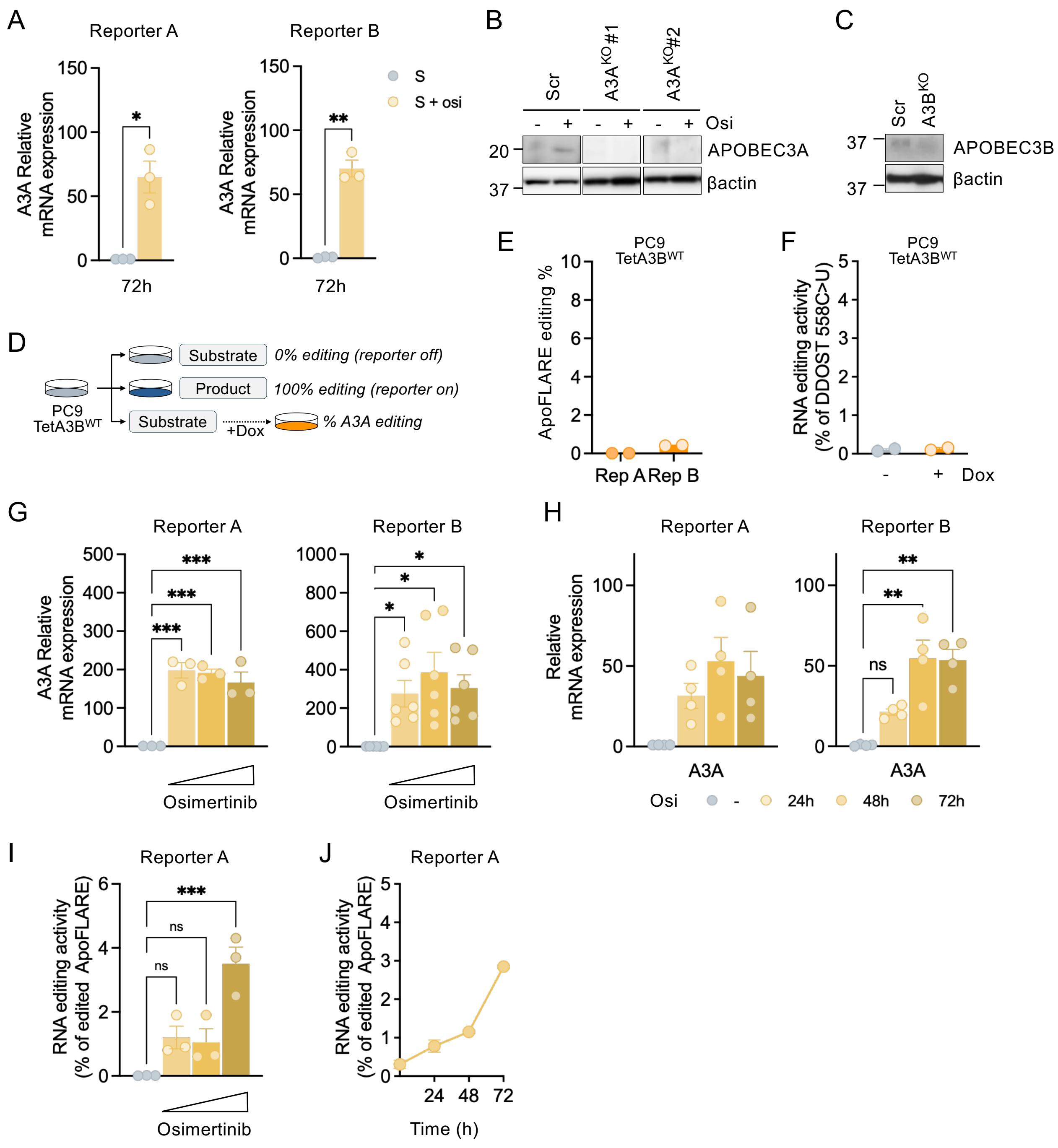


**Supplementary figure 4. A.** A3A expression in PC9 cells treated with osimertinib for 72 hours (mean ± SEM of three biological replicates, Student's t test). **B, C**. Immunoblot of A3A (**B**) or A3B (**C**) expression following genetic knockout after CRISPR-RNP delivery. $A3A^{KO}$ cells were treated with vehicle or osimertinib. β-actin was used as a loading control. **D**. Experimental schema for testing ApoFLARE in PC9 cells with inducible A3B. **E, F**. ApoFLARE editing (Reporter A and B) in PC9 $TetA3B^{WT}$ cell lines treated with dox for 72 hours, and corresponding DDOST 558 C>U mRNA editing determined by ddPCR. (mean ± SEM of two biological replicates). **G, H**. A3A expression in PC9 cells treated with increasing concentrations of osimertinib (0, 10 nM, 100 nM, 1 μM) for 72 hours (**G**) or osimertinib (300 nM) for the indicated timepoints (**H**) (mean ± SEM of three to six biological replicates, one-way ANOVA followed by Sidak's post-hoc test). **I, J**. ApoFLARE reporter A mRNA editing determined by allele-specific ddPCR. (mean ± SEM of two to three biological replicates; **I**, one-way ANOVA followed by Sidak's post-hoc test). Dox, doxycycline; osi, osimertinib.
